# Genetic heterogeneity in autism: from single gene to a pathway perspective

**DOI:** 10.1101/042218

**Authors:** Joon Yong An, Charles Claudianos

## Abstract

The extreme genetic heterogeneity of autism spectrum disorder (ASD) represents a major challenge. Recent advances in genetic screening and systems biology approaches have extended our knowledge of the genetic etiology of ASD. In this review, we discuss the paradigm shift from a single gene causation model to pathway perturbation model as a guide to better understand the pathophysiology of ASD. We discuss recent genetic findings obtained through next-generation sequencing (NGS) and examine various integrative analyses using systems biology and complex networks approaches that identify convergent patterns of genetic elements associated with ASD. This review provides a summary of the genetic findings of family-based genome screening studies.

## 1. Introduction

Autism spectrum disorder (ASD) is a highly heritable neurodevelopmental disorder that affects up to 1-2% of the population (Autism et al., 2012; Devlin and Scherer, 2012). It is characterized by impaired social interactions, communication deficits and repetitive behavior. Moreover, ASD is often comorbid with other phenotypic and clinical features, such as motor impairments, sleep disturbance, and sensory abnormalities. Not surprisingly, ASD is characterized by both genetic and phenotypic heterogeneity, and there is a high level of variation between individuals diagnosed with autism.

Over the last five years, technological advances have facilitated identification of causal DNA variants associated with ASD. Advances in next-generation sequencing (NGS), whole-exome sequencing (WES), and whole-genome sequencing (WGS) have enabled the identification of large numbers of rare single nucleotide variants (SNVs), including small insertion/deletion mutations (indels) that are associated with ASD. Furthermore, family-based sequencing (trio and sibling families) has helped to disentangle the causal relationship of *de novo* and inherited variants (Ku et al., 2013; Stein et al., 2013), such that, we now understand there are hundreds of genetic variations that affect a wide range of molecular functions associated with ASD (Betancur, 2011a; Buxbaum et al., 2012; Stein et al., 2013). More importantly, these genes are not randomly distributed but converge in functionally relevant biological processes such as synapse development and transcriptional regulation in the brain. Thus, the challenge that arises is to find and precisely identify potentially hundreds of heterogeneous DNA variations that converge in functional pathways associated with ASD. This requires refocusing molecular related morbidity paradigms from single or a few unrelated coding genes to functional networks of genes that contribute to key neurodevelopmental and neurological processes.

Systems biology and complex networks approaches are now being used to build the relationships between biological elements among heterogeneous components. This has been important to focus on a level of resolution that allows us to ‘see the forest for the trees’. We are mindful that ‘nothing in biology makes sense except in the light of evolution’ (Dobzhansky, 1973), and that evolutionary constraint underpins the fidelity of genes and molecules that function in biological process. Importantly, this has led to a shift from an exclusive reliance on statistical genetics to more appropriately understanding the evolutionary preconfigured nature of a biological system and complex molecular networks that underpin human development. The application of the systems paradigm in mental health research allows us to examine the relevance of combinations of genetic elements that putatively impact on biological functions associated with ASD.

Much of the scientific momentum regarding systems approaches comes from traction arising from many different omics data (Mitra et al., 2013), increasing our capacity to generate new models, for hypothesis-driven analyses of genetic data (Barabasi et al., 2011). This has led to the development of integrative approaches involving multiple data sources, model construction, and complex network analyses. Here, we provide an overview of recent WES and WGS studies for large ASD cohorts. We are mindful, the systems paradigm does not advocate abandoning statistical approaches but rather refines the search space for probability analysis to be appropriately applied. This review summarizes various integrative approaches that were made possible by advancements in systems biology methods.

## 2. Recent genetic findings in ASD

### 2.1. *De novo* variants

A major genetic finding was the identification of *de novo* variants (DNVs) associated with ASD. DNVs are DNA variants that occur spontaneously in the parental germline during meiosis (Conrad et al., 2011). Similar to inherited variants, DNVs comprise a wide range of genomic alterations, including chromosomal variations, copy-number variations (CNVs; >1kb), small insertions/deletions (indels; 2-1,000bp) and single nucleotide polymorphisms (SNPs; 1bp). On average, approximately 40 DNVs are thought to arise in an individual’s genome (Conrad et al., 2011). Most DNVs appear to be correlated with paternal age (Flatscher-Bader et al., 2011; Hultman et al., 2011; Malaspina et al., 2001; McGrath et al., 2014), as aging germ-line cells involved in spermatogenesis introduce errors during replication, thereby increasing the rate of DNVs (Kong et al., 2012; O’Roak et al., 2011). In sporadic cases of ASD (i.e., ASD probands with unaffected parents), DNVs have been reported to have a large genetic effect, as they have not been subject to natural selection processes in previous generations (Crow, 2000; Eyre-Walker and Keightley, 2007).

A number of previous large-scale studies focused on copy number variations (CNVs) using SNP genotyping arrays and identified *de novo* events associated with ASD (Levy et al., 2011; Marshall et al., 2008; Pinto et al., 2010; Sanders et al., 2011; Sebat et al., 2007; Szatmari et al., 2007). The results highlight several loci where recurrent *de novo* CNVs, including 1q21.1, 7q11.23, 15q11-13, 16p11.2, 17q12, and 22q11.2. These CNVs are larger duplications or deletions that encompass multiple genes or regulatory regions and are expected to impart a substantial risk for certain phenotypes. Many of these *de novo* CNVs were shown to affect important synaptic genes, such as *NRXN1* (Szatmari et al., 2007), *NLGN3* (Sanders et al., 2011), *SHANK3* (Marshall et al., 2008), *SHANK2* (Berkel et al., 2010), and *SYNGAP1* (Pinto et al., 2010), suggesting synaptic dysfunction or perturbation of synapse development may underlie the pathophysiology of ASD. Indeed, subsequent studies using animal models and transcriptomics data have elucidated functional changes associated with these CNVs (Durand et al., 2012; Golzio et al., 2012; Luo et al., 2012; Won et al., 2012). *De novo* CNVs have an increased frequency in individuals with ASD compared with unaffected controls, in contrast to inherited CNVs, which occur at similar rates between cases and controls. The enriched occurrence of *de novo* CNVs in ASD cases (accounting for only 5-10% of total ASD cases) and low co-occurrence in unaffected controls (~1%) (Carter and Scherer, 2013; Geschwind, 2011) indicate inherited CNVs are often not causally related and that a minor percentage of *de novo* CNVs are not fully penetrant in ASD (Bucan et al., 2009).

Despite some success showing CNVs being causatively associated with ASD, the SNP genotyping array approach is restricted by the large genomic intervals between SNP markers and cannot be used to reliably investigate SNPs or indels that occur in this intervening space. In contrast recent advances using massive-throughput genome sequencing, allows point mutations and indels (DNA variations) to be examined on the basis of hundreds of millions of sequence reads. For example, high-fidelity whole-exome sequencing (WES) focuses on genetic information of protein-coding transcripts and their nearby regulatory regions (Bamshad et al., 2011). Alternatively, whole-genome sequencing (WGS) provides a more comprehensive readout across the entire genome but it is computationally difficult to analyze and currently still significantly more expensive than WES.

Since the introduction of WES and WGS, increasing evidence has linked DNVs with ASD. Recent family-based studies identified thousands of DNVs in a large number of ASD probands and their siblings (Table 1). Loss-of-function variants (LoF variants) – those annotated as nonsense variants, frameshifts or splice site changes – occur at twice the frequency in probands (roughly 0.21 per child) than in unaffected siblings (roughly 0.12 per child) (Iossifov et al., 2014). For this reason, *de novo* LoF variants are considered to provide one of the major genetic contributions to sporadic cases (Hoischen et al., 2014; Ronemus et al., 2014). Importantly, two recent large exome studies, involving 3,871 and 2,508 cases revealed multiple *de novo* LoF variants occur in coding genes – *ADNP*, *ANK2*, *ARID1B*, *CHD8*, *DYRK1A*, *GRIN2B*, *KATNAL2*, *POGZ*, *SCN2A*, and *TBR1* (Iossifov et al., 2014; Poultney et al., 2014).

**Table 1.** *De novo* variants identified through exome and genome sequencing

| Study | Method | Probands | Siblings |  | LoF* | Missense** | Others*** | Total |  | LoF* | Missense** | Others*** | Total |
| --- | --- | --- | --- | --- | --- | --- | --- | --- | --- | --- | --- | --- | --- |
| O'Roak 2012 | WES | 209 | 50 | DNVs in probands | 38 | 154(92) | 68 | 260 | DNVs in siblings | 3 | 31(21) | 16 | 50 |
| Sanders 2012 | WES | 225 | 200 |  | 15 | 125(87) | 32 | 172 |  | 5 | 82(54) | 38 | 125 |
| Neale 2012 | WES | 175 | 0 |  | 18 | 101(NA) | 50 | 169 |  | NA | NA | NA | NA |
| Iossifov 2012 | WES | 343 | 343 |  | 61 | 209(NA) | 92 | 362 |  | 30 | 203(NA) | 81 | 314 |
| Jiang 2013 | WGS | 32 | 0 |  | 5 | 28(13) | 9 | 42 |  | NA | NA | NA | NA |
| An 2014 | WES | 48 | 0 |  | 3 | 18(11) | 8 | 29 |  | NA | NA | NA | NA |
| De Rubeis 2014 | WES | 3,871 | NA |  | 307 | 2,186 (828) | NA | 1,135 |  | NA | NA | NA | NA |
| Iossifov 2014 | WES | 2,508 | 1,911 |  | 383 | 1,667 (NA) | 646 | 2,696 |  | 178 | 1,140 (NA) | 490 | 1,808 |

\* LoF variants are nonsense variants, frameshifts or splice-site-changing variants
\*\*The numbers in brackets indicate the number of missense variants classified as putatively deleterious by functional prediction algorithms (e.g., SIFT, PolyPhen2).
\*\*\* Including synonymous and non-frameshift variants

*De novo* missense variants are the most abundant type of DNV (approximately 60%) identified in WES and WGS studies. Although they are thought to account for approximately 10% of ASD cases (Ronemus et al., 2014), it is difficult to determine whether *de novo* missense variants show greater frequency in ASD probands than in unaffected siblings. Recent studies reported the rate of *de novo* missense variants was increased in probands versus siblings (O’Roak et al., 2012; Sanders et al., 2012), but another study did not detect this difference (Iossifov et al., 2012). Nevertheless, there are a few factors that should be taken into consideration when estimating their contributions. Firstly, most *de novo* missense variants result in deleterious amino acid substitutions (Table 1), and their individual effects must be considered in terms of structural modification and functional consequence (binding, dimerization, catalysis, post-translational modification) to the protein or isoform protein. Secondly, a large number of *de novo* missense variants found in the same gene are required to establish a significant association with ASD (Sanders et al., 2012). Thirdly, *de novo* missense variants have been found to be overrepresented during early neurodevelopment and prenatal development (Parikshak et al., 2013).

Interestingly, there are numerous recurring variants observed in ASD probands. These include various types of DNVs that occur in coding regions of SCN2A (Sodium Channel, Voltage-Gated, Type II, Alpha subunit) were found in four different studies (An et al., 2014; Iossifov et al., 2012; Jiang et al., 2013; Sanders et al., 2012). We also know many of the recurrent DNVs have been associated with other neurodevelopmental disorders, such as intellectual disabilities, epileptic encephalopathy’s and schizophrenia, suggesting there is high molecular comorbidity between these disorders (Iossifov et al., 2014; Li et al., 2015; Poultney et al., 2014). Cristino *et al.* recently demonstrated there is significant molecular overlap between neurodevelopmental and neuropsychiatric disorders (Cristino et al., 2014a). Apart from recurrent genes containing DNVs, it is noteworthy that there is an increased frequency of DNVs occurring in several gene families. These include voltage-gated calcium channel (*CACNA*), catenin (*CTNN*), and chromodomain helicase DNA binding protein (*CHD*) gene families that appear to contribute to specific ASD phenotypes (Bernier et al., 2014; Pinggera et al., 2015; Turner et al., 2015). This increased frequency of DNVs in specific gene families highlights related genes with overlapping function contribute to biological pathways and processes associated with synapse development and function, neuronal development, and chromatin modulation (Figure 1).

**Figure 1.**
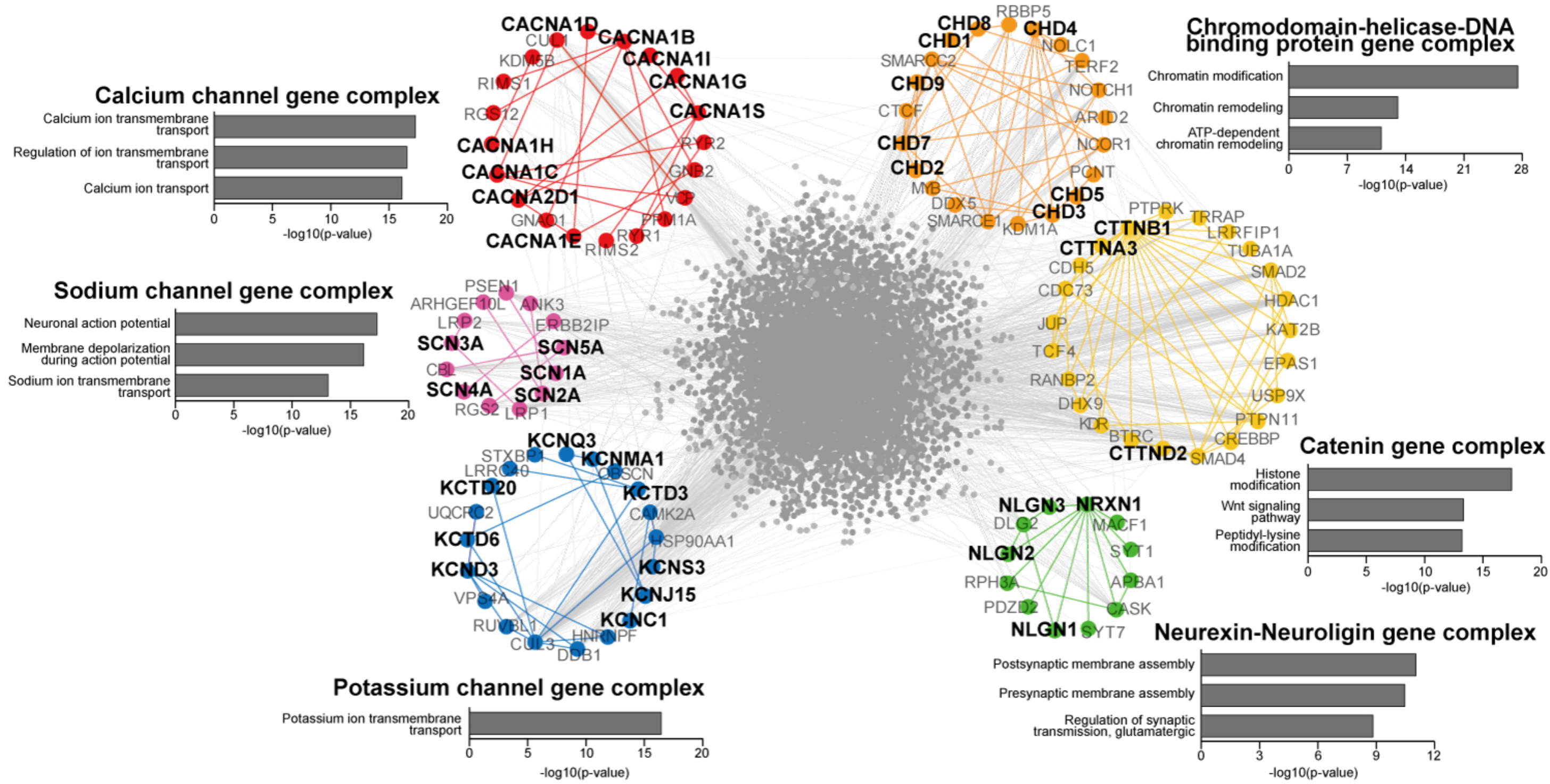
Molecular pathways of *de novo* variants (DNVs) associated with ASD. The protein-protein interaction network plot illustrates six gene families that DNVs were frequently occurred in ASD cases. Interactions were retrieved from HPRD and BioGrid. The sub-network contains DNVs of gene complex and their first-degree interactors. Gene set enrichment analysis was performed to the sub-network genes using the Gene-Ontology database. Bar graphs show top significant pathways associated with the sub-network genes. The y-axis in the bar graph is an enrichment score, provided in a log10-transformed p-value.

### 2.2. Rare inherited variants

In contrast to DNVs, inherited variants account for the greatest number of genetic alterations in human genomes and can be classified as either common or rare on the basis of their frequency in the population. Although both common and rare inherited variants may contribute to the development of ASD, rare variants are expected to have a larger effect than individual common variants (State and Levitt, 2011; Stein et al., 2013). Importantly, recent, WES studies have successfully identified a range of rare inherited variants associated with ASD (An et al., 2014; Lim et al., 2013; Toma et al., 2013; Yu et al., 2013).

Rare complete knockouts (homozygous and compound variants) have typically been associated with Mendelian disorders because they more obviously result in deleterious phenotypes. Lim *et al.* recently reported rare complete knockouts in ASD cases using WES analysis of 933 ASD cases and 869 controls to examine (Lim et al., 2013). Among the 933 ASD cases and 869 controls examined there were 91 rare complete knockouts were found. These occur at twice the frequency in cases (n=62) than in controls (n=29), and the authors concluded this incidence accounts for ~3% of ASD cases. This disparity was not observed for rare heterozygous variants. Of particular interest was the high occurrence of rare complete knockouts associated with the X-chromosome in cases compared with the controls. The predominant of hemizygous variants found in males or homozygous variants in females are estimated to account for another 2% of ASD cases. In another study by Yu *et al.*, heterozygous mapping and linkage analysis was used to examine consanguinity in familial ASD cases (Yu et al., 2013). The authors found rare LoF and missense variants associated with *AMT*, *PEX7* and *SYNE1* genes. They also examined protein-altering variants (missense, nonsense, splice site, and frameshift mutations) present at low frequencies in the population. The analysis of 70 genes previously associated with ASD identified an excessive number of monogenic, autosomal recessive or X-linked variants (Betancur, 2011b). They found five families affected by rare homozygous, compound heterozygous, or hemizygous variants. For example, a nonsense hemizygous variant occurring in *NLGN4X* was identified in one family; this variant was inherited from an unaffected mother in an affected male child. Another family showed the same inheritance pattern for a nonsense hemizygous variant of *MECP2*. Although some rare homozygous variants can explain the etiology of some families with clinical phenotypes associated with certain syndromes (e.g., phenylketonuria, dysmorphic features or microcephaly) others occur in unaffected siblings or cases that do not show clear phenotypes, suggesting hypomorphic variants or other genes might be involved.

Two WES studies have analyzed the role of rare heterozygous variants in ASD. In some multiplex ASD families rare heterozygous truncating variants – nonsense and frameshift variants – are more frequently inherited from parents than non-truncating variants. More importantly, there is an inverse correlation between the number of rare heterozygous truncating variants and non-verbal intelligence quotient (NVIQ) has been observed (Toma et al., 2013), confirming previous observation concerning an increased number of DNVs and decreased NVIQ (O’Roak et al., 2012). However, this study did not ascertain the contribution of rare heterozygous missense variants. More recent evidence showed that rare heterozygous missense variants appear to contribute to the etiology of ASD. Recently, An *et al.* (2014) investigated significance of rare heterozygous variants identified by stringent filtering of sequence data. Rare variants were defined as those that a) showed a minor allele frequency (MAF) of <1% or were unknown in the 1,000 Genomes Project and NHLBI Exomes database and b) were predicted to be deleterious by SIFT and PolyPhen2. This analysis identified ~125 rare variants per individual. These variants were significantly enriched in genes that were previously associated with ASD and correlated with the level of rarity (i.e. MAF or prediction scores). Moreover, inherited variants from parents with broader autism phenotype (BAP), a weak phenotype of ASD (Taylor et al., 2013), were enriched in ASD-associated genes compared with inherited variants from parents without BAP. Taken together, this evidence indicates that rare heterozygous variants make a cumulative contribution to ASD.

## 3. Convergent molecular aberrations underlying ASD

The extreme genetic heterogeneity found in ASD represents a huge challenge in attempting to understand the etiology and development of autism (Betancur, 2011b; Geschwind, 2011). In the absence of any specific gene responsible for the majority of ASD cases, the most common genes only account for approximately 2% of ASD (Abrahams and Geschwind, 2008). Furthermore, it has been shown that candidate genes cover a wide range of functions, including cell-cell adhesion, synaptic activities and neurotransmission, transcriptional regulation, and activity-dependent protein translation, and possibly contribute to a number of common clinical features that accompany ASD (Abrahams and Geschwind, 2008), which are generally devalued in the diagnostic processes and in the discovery of candidate genes (Jeste and Geschwind, 2014; Stessman et al., 2014). For this reason, ASD should be considered as a combination of phenotypes, not as a disorder caused by any individual gene (Geschwind, 2011; Mitchell, 2012). It is therefore essential to examine the contribution of the many candidate ASD genes with respect to functions that contribute to diagnostic behaviors in order to elucidate the numerous subtypes of ASD.

This situation has led to the emergence of new approaches for detecting molecular convergence associated with ASD. There are now a number of studies that have attempted to resolve whether putative candidate genes converge in specific molecular or biological processes. These studies are based on the premise that convergent pathway(s) and the co-expression or functional proximity of candidate genes display distinctive patterns of interaction and provide a *segue* for systems biology analyses.

### 3.1. Interconnection of ASD genes

Many studies have investigated the convergence of ASD-associated CNVs in biological pathways. Gilman *et al.* developed the Network-Based Analysis of Genetic associations (NETBAG) technique for this purpose (Gilman et al., 2011). This method was designed to identify combinations of genes contributing to any one disorder, based on the hypothesis that these genes contribute to phenotypes typical of a disorder. NETBAG used high-confidence datasets of ASD-associated CNVs (Levy et al., 2011) and showed these were enriched in pathways related to synaptogenesis, cell-cell adhesion, axon guidance, neuronal motility, GTPase signaling and the actin cytoskeleton. This method was later applied to examine a large sample set, and helped to identify a network of 113 *de novo* CNVs, enriched in functions associated with chromatin and transcription regulation, *MAPK* signaling, and synaptic development (Pinto et al., 2014). Interestingly, putative functional perturbations associated with CNVs are more pronounce in females than in males, consistent with the idea of highly penetrance CNVs occurring in females (Pinto et al., 2014). By considering duplication and deletion variations, Noh et al. have further elucidated the phenotypic consequences of CNVs (Noh et al., 2013).

Importantly, pathways relevant to synaptic functions have repeatedly been associated with ASD pathology (Betancur et al., 2009; Toro et al., 2010). Sakai *et al.* (Sakai et al., 2011) highlight how CNVs converge in neuronal post-synaptic processes in ASD. Specifically, they used a yeast two-hybrid system to examine protein-protein interactions. They found genes from ASD cases had greater connectivity compared to non-ASD individuals. Many key genes (or hub genes) that interact with a number of other genes (protein-protein interactions) are more frequently associated with CNVs found in ASD cases. For example, *SHANK3* is connected with a number of postsynaptic genes (e.g., *PSD95*, *TSC1*, *ACTN1*, and *HOMER3*). *In vivo* evidence (Becker et al., 2014; Tsai et al., 2012) confirms the post-synaptic complex is an important contributor to ASD pathophysiology (Berg and Geschwind, 2012).

This integrative approach provides a basis for constructing more comprehensive models for conceptualizing the molecular relationship between ASD and other neurodevelopmental and neuropsychiatric disorders. Cristino *et al.* (2014a) examined the functional patterns of candidate genes associated with ASD, X-linked intellectual disability (XLID), attention deficit hyperactivity disorder (ADHD) and schizophrenia (SZ). In this study, candidate disorder genes and their first-degree interacting proteins were used to create a comprehensive protein-protein interaction network, which was collectively referred to as the AXAS-PPI network. Interestingly, the AXAS-PPI network was shown to have significantly more connections with increased structural properties (e.g., in terms of the average degree, density and clustering coefficient) compared to networks created from an equivalent number of randomly selected genes in the genome. This confirmed the AXAS network reflects meaningful biological signal, confirming neighboring candidate genes function in the related biological processes and pathways. Closer examination revealed this AXAS network is divided into 30 functional modules, with unique patterns of gene distributions across the four disorders and this can be used to analyze the relationship of DNA variations with different disorders. For instance, the module for synapse development and signal transduction was shown to be the largest with disorder genes being equally distributed. Synapse development and signal transduction may therefore be key biological processes common to ASD, XLID, ADHD and SZ. This is consistent with the number of studies that highlight the importance of synaptic genes (*DLG3, DLG4, GRIN1, GRIN2B, PRKCB, STX1B, STXBP1*, and *SYNGAP1*) in ASD, epilepsy, intellectual disabilities and schizophrenia (Hormozdiari et al., 2015).

Complex network models are now proving to be powerful tools for analyzing genetic screening data and abstracting the molecular basis of neurodevelopmental and neuropsychiatric disorders. An *et al.* (2014) utilized the AXAS model to analyze WES sequencing data and identify patterns of candidate genes that are putatively overrepresented in functional processes and pathways associated with ASD. They found approximately 100 rare variants per individual case were distributed across a number of different biological pathways and processes. For instance, rare variants from four ASD probands were clustered in the *L1CAM* pathway including combinations of rare inherited variants and DNVs. This result showed there was molecular convergence of predominantly heterozygous variants in families with ASD. Their data suggests combinations of weak heterozygous alleles converge in common biological processes and likely contribute to increase threshold of liability associated with ASD. Again, the clustering phenomena were observed in pathways related to synaptic function (e.g. the neurexin trans-synaptic complex). Indeed, by examining the distribution of rare variants among interacting proteins involved in synapse development and function, they confirm previously reported pattern of pervasive rare variants (Kenny et al., 2013).

The interconnection of ASD genes has been further investigated by considering the expression of alternative spliced genes. Corominas *et al.* (2014) built a PPI network, the Autism Splice-form Interaction Network (ASIN), based on alternatively spliced brain-expressed isoforms and genetic associations (Corominas et al., 2014). The background interactions of the ASIN consisted of experimentally validated isoforms generated from high-throughput transcriptomics sequencing, but the data suffered from a lack of spatiotemporal evidence due to the use of only two adult brain samples (one from an 18-year-old male and another from a 66-year-old female). Interestingly, the interacting partners of the ASIN were enriched for rare *de novo* CNVs. This is consistent with findings in other network-based studies (Cristino et al., 2014a; Liu et al., 2014), and implicates functional proximity as a potential measure to identify other candidate ASD genes. For instance, *de novo* LoF variants in GATAD2B have been observed in patients with limited speech (de Ligt et al., 2012), and due to its physical interaction with FOXP2, impairment of GATAD2B may explain this language development issue. Another study proposed a composite gene model by integrating brain-specific exon expression and allele frequencies in the general population (Uddin et al., 2014). This model assumed the integrity of gene expression to be modulated by selective pressure on particular isoform proteins, thus ranking the importance of genes that confer susceptibility. In this model, genes showing high exonic expression in the brain and a low allele frequency significantly overlapped with known ASD candidate genes relevant to fragile-X protein targets.

### 3.2. Transcriptional convergence on ASD and neurodevelopment

Co-expression networks are widely used for investigating the relationship between genes in specific spatial and temporal contexts. These networks are emerging as a powerful tool and incorporate recent high-resolution transcriptional profiles associated with human brain development (Hawrylycz et al., 2012; Kang et al., 2011; Miller et al., 2014).

The most common approach to look at transcriptome data is Weighted-Gene Co-expression Network Analysis (WGCNA) (Langfelder and Horvath, 2008). WGCNA has been used to survey modules of highly correlated genes typical of specific brain regions and developmental stages. Voineagu *et al.* analyzed postmortem brains to demonstrate common pathophysiological abnormalities associated with cortical areas in ASD cases (Voineagu et al., 2011). They again substantiated numerous synaptic and immune genes were found to be perturbed in the frontal and temporal cortex of ASD cases. WGCNA was further applied to examine the global differences in the organization of the brain transcriptome. This analysis revealed salient differences in regional gene expression between autism cases and controls. In the frontal and temporal cortex, gene expression patterns varied significantly among the controls but not in ASD cases, suggesting there are specific regional abnormalities in the brain. Furthermore, co-expression modules similarly confirm synaptic genes were upregulated, whereas immune-related genes were downregulated. The synaptic gene module was found to be enriched with genome-wide association signals, subsequently confirmed by others (Ben-David and Shifman, 2012). Similarly, in a recent study by Gupta *et al.* (2014), the authors used WGCNA to identify the negative correlation between the module of activated M2-state microglia genes and that of synaptic transmission genes from 47 ASD postmortem brains (Gupta et al., 2014).

Several studies have now combined DNA variations discovered from genome sequencing with co-expression network analysis to demonstrate there is a convergence of genetic variations in the perinatal/postnatal cerebellar cortex and layer 5/6 cortical projection neurons (Parikshak et al., 2013; Willsey et al., 2013). Parikshak *et al.* (2013) also demonstrated ASD candidate genes associated with the superficial cortical layers during early neurodevelopment. The analysis of co-expression networks found that DNVs in ASD cases were enriched in the transcriptional regulation module but common inherited variants were enriched in the synaptic transmission module. Furthermore, co-expression modules highlighted the temporal and spatial difference contributes to developmental trajectories. From the prenatal period to infancy, the level of genes involved in transcriptional regulation decreased, whereas the levels of genes involved in synaptic transmission increased. In the adult cortex, rare *de novo* and common inherited variants were clustered in superficial cortical layers 2–4 and upper layer glutamatergic neurons. A study by Willsey *et al.* (2013) demonstrated the gene expression changes associated with *de novo* LoF variants in deep projection neurons of the mid-fetal frontal cortex. Furthermore, these variants were enriched at the inner cortical plate and the cortical glutamatergic neurons. Similarly, Li et al. reported cortical disconnections in ASD cases (Li et al., 2014). Combining genomic and RNA sequencing data, they found rare nonsynonymous variants occur in genes involved in oligodendrocyte development, which is the major cell type of the corpus callosum. Taken together, these observations suggest genetically-driven developmental *diaschisis* occurs, whereby early dysfunction may disrupt the maturation of cortical circuits in the brain and contribute to ASD phenotypes (Wang et al., 2014).

The above examples indicate co-expression network analysis can be integrated into a risk prediction model for ASD. The Detecting Association With Networks (DAWN) model proposed by Liu *et al.* (Liu et al., 2014) determined ‘hot spots’ of genetic associations within a co-expression network. In addition to the statistical assessment of *de novo* and inherited variants (He et al., 2013) showed DAWN was capable of prioritizing possible high-risk genes in the mid-fetal pre-frontal and motor-somatosensory neocortex. This approach was able to provide a rigorous assessment and identification of causal genes that includes evidence from targeted sequencing. Another feature of this model was the identification of sub-networks centered on *FOXP1* and *PTEN* clearly demonstrate convergence of ASD-associated genes in neural organization and language development.

## 4. Concluding remarks

In this review, we provide an overview of published ASD studies based on genome screening and systems biology. Recent advances using pedigree-based genome screening have enabled more rigorous examination of genetic variations. WES and WGS studies have led to the discovery of rare *de novo* and inherited variants with putative greater penetrance than predicted common variants. Moreover, there is great utility with these approaches to interpret biological significance of genetic variations 1) examining causative DNA variations based on an actual nucleotide position (Ku et al., 2013), 2) estimating effect of DNA variations on gene expression in key pathways (Deriziotis et al., 2014), 3) utilizing information of minor allele frequency and evolutionary conservation to characterize rare variations associated with ASD (An et al., 2014).

It is now clear that there is no single ASD-causing gene. Intensive research over the last five years has confirmed there is an extensive genetic heterogeneity associated with ASD. This is supported by understanding that DNA variations often converge in biological pathways and functional processes and are shared across a broad range of neurodevelopmental and neuropsychiatric disorders or specific to only one disorder (Sullivan and Posthuma, 2015). Therefore, systems biology approaches will be a valuable tool to dissect specific differences between disorders (Cristino et al., 2014a; Parikshak et al., 2013; Willsey et al., 2013), and identify functional distribution of causal DNA variants in individuals with ASD (An et al., 2014).

There are now an impressive range of technologies that can be used investigate functional consequences of genetic variations. Induced pluripotent stem cells (iPSCs) are currently being used to characterize the effect of genetic variations in neuronal cell lineages involved in early neurodevelopment (Griesi-Oliveira et al., 2014). Generating human neurons from iPSCs has potential utility comparing healthy and affected individuals that will help characterize the role of causal ASD genes. The use of clustered regularly interspaced short palindromic repeats (CRISPR)/Cas9 gene editing system with iPS cells provides a tractable method for *in vitro* analysis of DNA variations whereby a number of overlaid genetic elements can be examined additively or in a subtractive fashion to assess relative contribution to risk genes to ASD phenotypes (Bernier et al., 2014). Similarly, these gene editing technologies can be used *in vivo* and aside from rodent models, high throughput small model species (zebra fish, fly, honeybee and worm) are now being used to validate the biological relevance of conserved genes and human genetic variations (Burne et al., 2011), including their contributions to complex behaviors, such as learning and memory (Cristino et al., 2014b).

In conclusion, systems biology approaches have facilitated an important paradigm shift from single gene to pathway etiology associated with ASD. Nevertheless, we are mindful of the enormous task of analyzing non-coding regulatory information with respect to DNA variations. Determining the functional impact of coding and non-coding regulatory DNA variations remains a significant hurdle for developing a truly integrated omic systems approach. Resources, such as ENCODE (Encode Project Consortium, 2012), FANTOM (Fantom Consortium, 2014), and mirBase (Kozomara and Griffiths-Jones, 2011), need to be integrated into current analysis pipelines to achieve a comprehensive predictive systems model for ASD.

## Acknowledgement

We thank Sarah Williams and Aoife Larkin for critical reading of the manuscript. J-Y An was supported by a University of Queensland PhD scholarship. C.C. was supported by funding from the Australian Research Council (FT110100292), National Health and Medical Research Council (APP1008125).

---

**Table 2.**
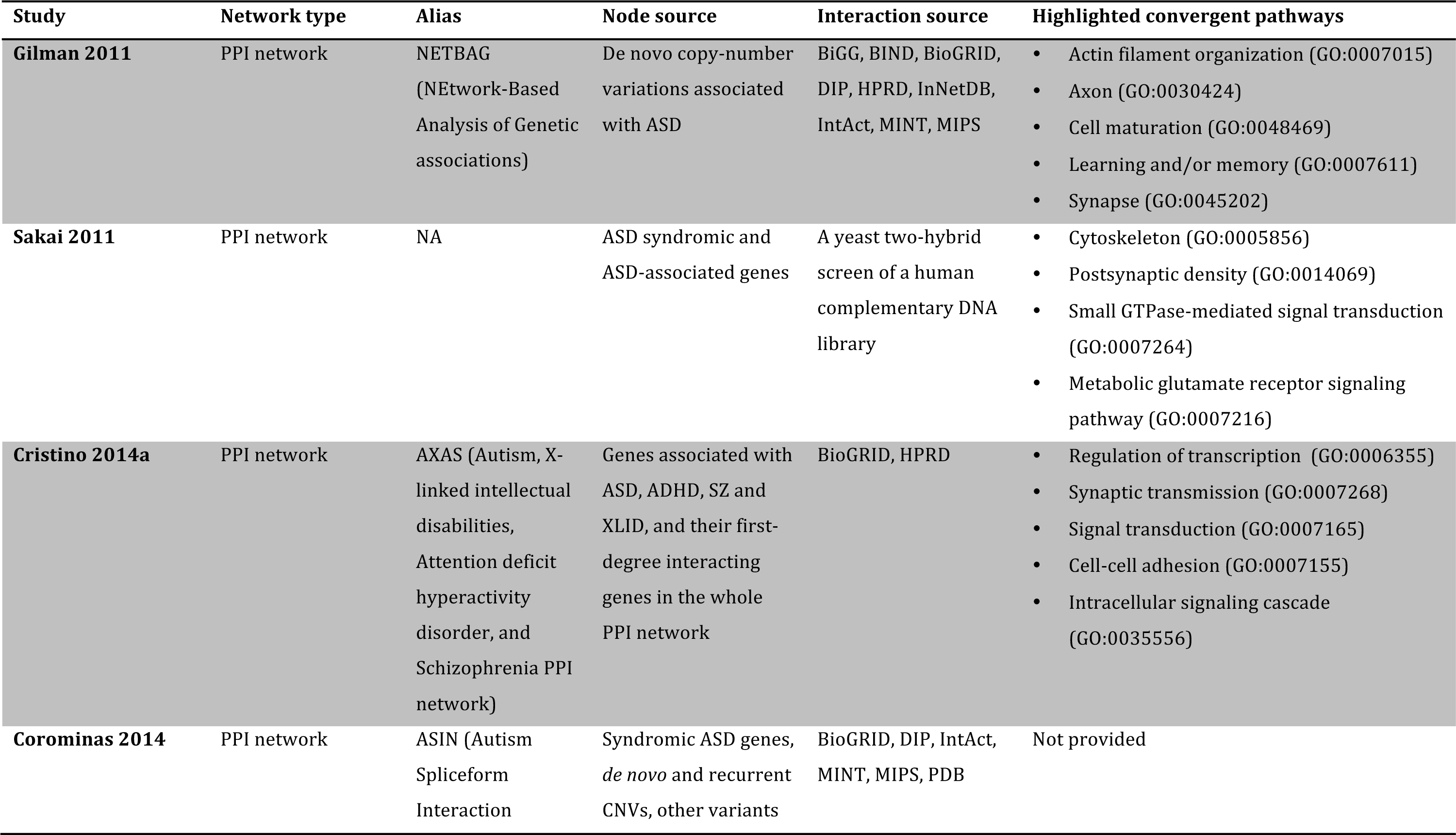
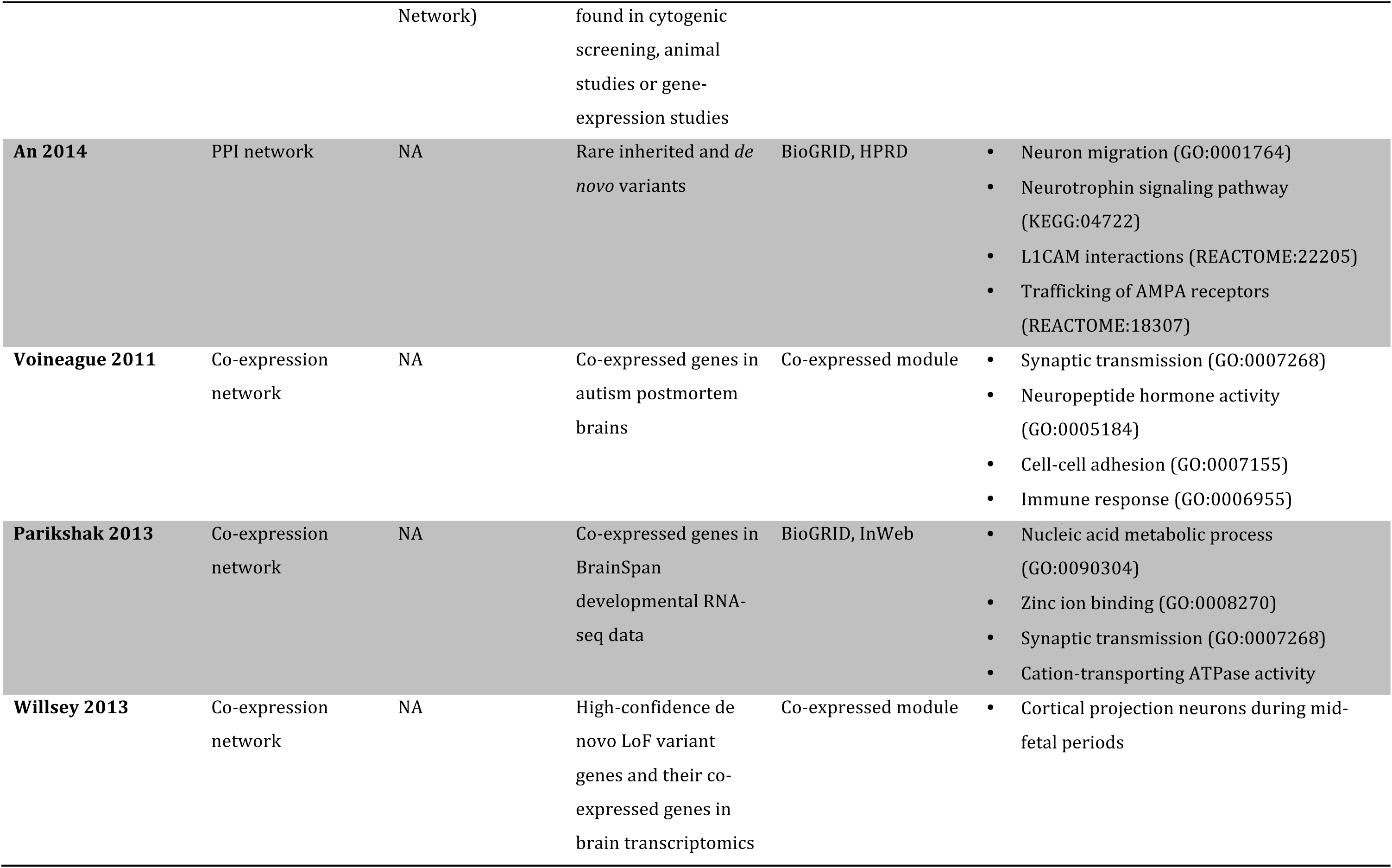
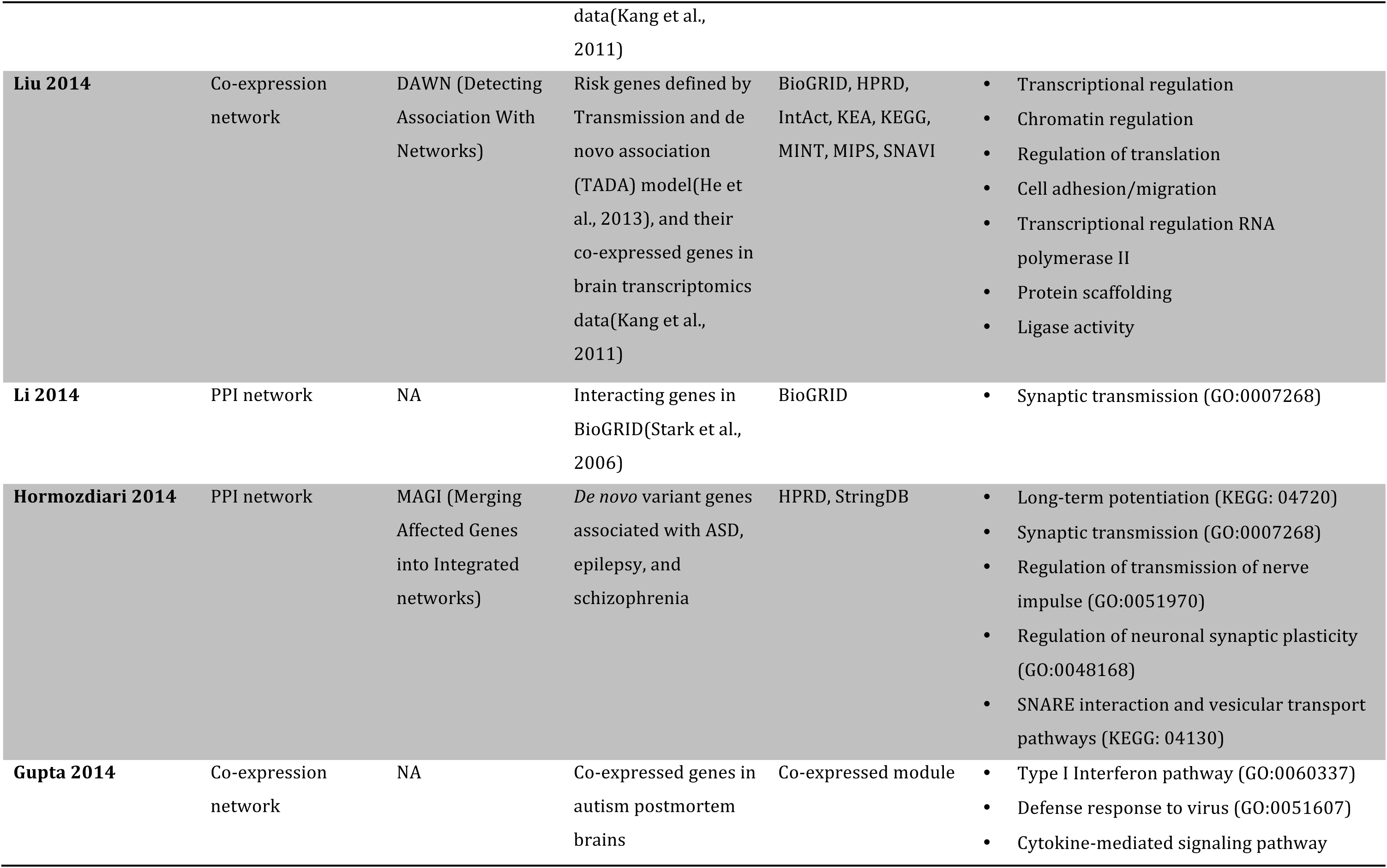
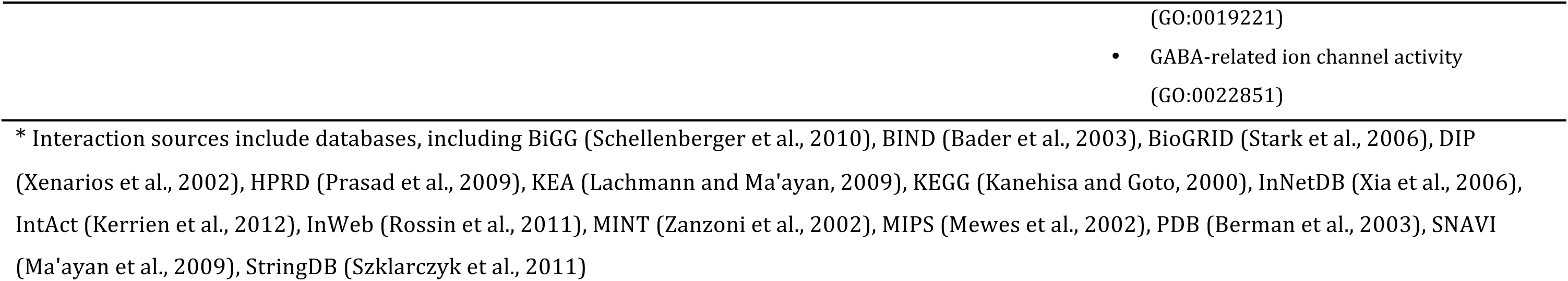
Network approaches in ASD studies

